# Gfi1-expressing Paneth cells revert to stem cells following intestinal injury

**DOI:** 10.1101/364133

**Authors:** Min-Shan Chen, Yuan-Hung Lo, Joann Butkus, Winnie Zou, Yu-Jung Tseng, Hsin-I Jen, Shreena Patel, Andrew Groves, Mary Estes, Ergun Sahin, Mark Frey, Peter Dempsey, Noah Shroyer

## Abstract

**Background＆Aim:** Chemotherapy drugs harm rapidly dividing normal healthy cells such as those lining the gastrointestinal tract, causing morbidity and mortality that complicates medical treatment modalities. Growth Factor-Independent 1 (GFI1) is a zinc finger transcriptional repressor implicated in the differentiation of secretory precursors into Paneth and goblet cells in the intestinal epithelium. We hypothesize that stimulating the reversion of Gfi1^+^ secretory cells into stem cells will improve intestinal epithelial regeneration and mitigate injury.

**Methods:** Gfi1 reporter mice (*Gfi1^cre/+;^ ROSA26 ^LSL-YFP^*) were treated with Doxorubicin, radiation, anti-CD3 antibody, and rotavirus to induce intestinal injury. Mice and intestinal organoids (enteroids) were used to investigate cellular repair mechanisms following injury.

**Results:** Under homeostatic conditions, Gfi1-lineage cells are Paneth and goblet cells, which were non-proliferative and not part of the stem cell pool. After injury, Gfi1^+^ secretory cells can re-enter the cell cycle and give rise to all cell lineages of the intestinal epithelium including stem cells. Reversion of Gfi1-lineage cells was observed in other injury model systems, including irradiation and anti-CD3 treatment, but not in ISC-sparing rotavirus infection. Our results also demonstrated that PI3kinase/AKT activation improved cell survival, and elevated WNT signaling increased the efficiency of Gfi1^+^ cell reversion upon injury.

**Conclusions:** These findings indicate that Gfi1^+^ secretory cells display plasticity and reacquire stemness following severe damage. Moreover, PI3kinase/AKT and WNT are key regulators involved in injury-induced regeneration. Our studies identified potential therapeutic intervention strategies to mitigate the adverse effects of chemotherapy-induced damage to normal tissues and improve the overall effectiveness of cancer chemotherapy.

## Introduction

Cytotoxic chemotherapy is a commonly used therapeutic approach to treat a wide range of cancers. Chemotherapy targets rapidly proliferating cells, both cancerous and normal cells as they undergo cell division, including intestinal stem cells in the gastrointestinal (GI) tract. Chemotherapy-induced GI injury is characterized by a rapid loss of stem cells, damage to the gastrointestinal epithelium, and activation of the inflammatory cascade ^1^. Doxorubicin is a topoisomerase II inhibitor which causes DNA damage during the cell cycle resulting in cell death; it is used to treat a range of malignancies, including: metastatic breast cancer, ovarian cancer, multiple myeloma, and AIDS-related Kaposi sarcoma ^2, 3^. However, Doxorubicin-induced toxicities commonly occur at sites within the GI tract and account for dose-limiting effects ^4^. Understanding the underlying mechanisms involved in regeneration of the intestinal epithelium will help to develop effective mitigators to reduce chemotherapy-associated drug toxicity and enhance tissue regeneration. Such insights may also provide novel avenues of therapeutic intervention in the setting of other GI injuries, including those caused by radiation, inflammation, or infection.

The intestinal epithelium is one of the most rapidly renewing tissues in adult mammals, where the epithelial cells are continuously renewed every 3–5 days ^5^. The epithelial integrity and rapid turnover are supported by intestinal stem cells (ISCs) located at the base of intestinal crypts. ISCs contribute to all intestinal lineages, including absorptive lineage cells: enterocytes and colonocytes; and secretory lineage cells: mucus-secreting goblet cells, hormone-secreting enteroendocrine cells, and antimicrobial peptide-secreting Paneth cells. Historically, two types of multipotent ISCs were identified. The first, known as crypt base columnar (CBC) cells, divide rapidly and are sensitive to radiation and chemotherapy injury. The second, termed +4 cells, marked by *Bmi1, mTert, HopX*, Sox9 and *Lrig1*, serve as quiescent ISCs (qISCs), and are slowly cycling under homeostatic conditions but can be induced to reconstitute the epithelium after injury^5–12^. Recent landmark studies suggest that long-lived secretory progenitors, differentiated enterocytes or enteroendocrine cells, reacquire stemness and replace the loss of Lgr5^+^ stem cells ^13–16^. These studies indicate that non-stem cells in the intestinal epithelium have developmental plasticity when subjected to tissue injury.

Gfi1 is a zinc-finger transcriptional repressor that was originally discovered in the hematopoietic system where it functions in the differentiation of hematopoietic lineages and adult hematopoietic stem cell (HSC) quiescence ^17–19^. Gfi1 is essential for several other cell types throughout the body, including Pancreatic acinar and centroacinar cells, inner ear hair cells, and pulmonary neuroendocrine cells ^20–22^. In the intestine, Gfi1 plays an important role in directing differentiation of secretory precursors into goblet and Paneth cells. Gfi1 knockout mice lack mature Paneth cells, have fewer goblet cells, and increased number of enteroendocrine cells ^23, 24^. It is still unclear whether committed secretory cells can serve as injury-induced ISCs following severe injury. Our data suggest that Gfi1^+^ secretory cells represent a unique cellular population that is distinct from active ISCs, *Dill1*^+^ secretory precursors, enterocyte precursors and +4 qISCs, thereby making them an excellent target for future therapeutic mitigators.

The Wnt signaling pathway is crucial for maintaining ISC hemostasis. R-spondins are Wnt agonist that are ligands of Lgr5, and stimulate canonical Wnt signaling to maintain ISCs, enhance crypt growth, and stimulate epithelial regeneration ^25–28^. R-spondins have also been implicated as clinically valuable chemoradioprotectors ^29, 30^. Phosphoinositide 3-kinase (PI3K)/AKT signaling is known to be essential for cell survival, proliferation, apoptosis, differentiation, and migration ^31^. The PI3K/AKT pathway and its downstream targets ("mammalian target of rapamycin," mTOR, and glycogen synthase kinase-3), play critical roles in the control and coordination of regeneration in many tissues ^32–34^. In homeostasis, adult intestinal crypt cell survival and proliferation are not mTOR dependent. However, following injuy by irradiation, crypt regeneration and ISC/progenitor maintenance require mTOR ^34^. We found that activation of PI3K/ AKT pathway combined with high R-spondin treatment shortly after chemotherapy-induced injury significantly enhanced cell survival and stem cell reversion, indicating the PI3K/Akt pathway and the Wnt pathway are important for the maintenance of ISCs and intestinal epithelium regeneration.

Our study demonstrates that mature Gfi1^+^ secretory cells reacquire plasticity upon tissue damage, and reveal potential mechanisms for intestinal regeneration and stem cell reversion following severe injury.

## Methods

### Animals

Lgr5-EGFP-IRES-creERT2 (Lgr5-GFP; stock No. 008875) and ROSA26-loxP-stop-loxP-rtTA-IRES-EGFP (*R26*^LSL-GFP^; Stock No. 005670) mice were purchased from the Jackson Laboratory. The Gfi1Cre knock-in mice ^35^ were generated by Dr. Lin Gan at the University of Rochester and were generously provided to us for this study. ROSA26-loxP-flanked STOP-EYFP (*R26^LSL-EYFP^*) mice (The Jackson Laboratory, Stock No. 007903) were a gift from Dr. Andrew Groves at Baylor College of Medicine. ROSA26-loxP-flanked STOP-TdTomato (*R26^LSL-TdTomato^*) mice (The Jackson Laboratory, Stock No. 007905) were a gift from Dr. Hui Zheng at Baylor College of Medicine. For doxorubicin treatment, 20 mg/kg doxorubicin (Sigma, D1515) was injected (i.p.) per mouse. For adenovirus-mediated modulation of Wnt signaling, 5 × 10^8^ PFU of Ad-Fc or Ad-Rspodin1 (Ad-Rspo1)^25^ was injected (i.v.) per mouse. For PTEN inhibitor treatment (SF1670), 5 mg/kg of PTEN inhibitor was injected (i.p.) per mouse. For irradiation injury studies, mice were given 10 Gy whole-body irradiation. For anti-CD3 Ab treatment, 50 μg of anti-CD3 antibody^36^ was injected (i.p.) per mouse. For rotavirus (RV) infection, RV was administered by oral gavage ^37^. The mouse experiments were approved by the Institutional Animal Care and Use Committee at Baylor College of Medicine.

### Immunofluorescent and immunohistochemical staining

Intestinal tissues were fixed with 4% paraformaldehyde, embedded in paraffin, sectioned (5 μm), and mounted on Superfrost Plus slides (ThermoFisher). Samples were blocked with 2% bovine serum albumin (BSA; VWR) and 5% either donkey or goat serum (Jackson immunoresearch), and then incubated with primary antibodies at 4°C for overnight. After washing three times with PBST (Phosphate buffered saline with 0.05% Tween-20), the samples were incubated with appropriate fluorescent dye- or HRP-conjugated secondary antibodies at RT for 1 hour. Slides were washed with PBST and mounted with mounting solution (Vector Laboratories, H-1000) for observation with a confocal microscope (Nikon A1-Rs). Primary and secondary antibodies and conditions are listed in Table 2.

### *In vitro* culture of intestinal crypts

Intestinal crypts were isolated from mouse duodenum as previously described ^38^, and cultured *in vitro*. These crypts were mixed with ice-cold Matrigel (Corning) and plated onto wells of a 24-well cell culture plate that was placed in the 37°C incubator for 30 minutes. 0.5 ml basel minigut media (Advanced DMEM/F12 (Invitrogen), 1% L-Glutamine (Invitrogen), 1% Pen/Strep, 10 mM Hepes, 1% N2 supplement, 2% B27) supplemented with 0.5 μg/ml rRspondin (R&D), 0.1 μg/ml rNoggin (R&D) and 25 ng/ml EGF (R&D) was added into wells. Recombinant growth factors could be substituted by R-spondin and Noggin conditioned media for cell maintenance. Recombinant growth factors could be substituted by Wnt3a, R-spondin and Noggin conditioned media. The completed media (basal minigut media with 10% Rspondin conditioned media, 10% Noggin conditioned media and 25ng/ml EGF) on which the cells were maintained was replaced every 2–4 days. Other groups developed R-spondin2-expressing^39^ and Noggin-expressing^40^ cells for conditioned media.

### *In vitro* injury culture in mouse enteroids

Mouse enteroids were broken down using a 30G insulin syringe and were plated with 50 μl matrigel/per well in 24-well plates. Basal minigut media with 50% Wnt-3a conditioned media, 10% Rspondin conditioned media, 10% Noggin conditioned media and 25 ng/ml EGF were added to the cells for overnight. Next day, the media was aspirated and 0.5 μg/mL Doxorubicin dissolved in completed media was added to the cultured enteroids. The cells were incubated at 37°C for 4 hours, after which the enteroids were washed 3 times with room temperature PBS. The cells were treated with complete minigut media or supplemented with either extra Rspondin (total 60% conditioned media), 10 *μ*m PTEN inhibitor (SF1670, Cellagen Technology), or 10 *μ*m AKT inhibitor MK2206. After 48 hours, the media was aspirated and complete minigut was added. Five days post-Doxorubicin treatment, the enteroids were analyzed under optical microscope. For Wnt3a conditioned media, a Wnt3a-expressing L-cell line is commercially available (ATCC, CRL-2647).

### Flow cytometry and cell sorting

Intestinal crypts were isolated as described above and dissociated with Dispase (Invitrogen) 1.2mg/ml at 37°C for 30 minutes. 2000 U/ml DNase (Roche) was added and the dissociated cells were passed through a 40 μm strainer. For quantification of Lgr5-GFP stem cells, the cells were washed with ice-cold PBS and stained with EpCAM-APC to identify epithelial cells, while Propidium iodide was utilized to determine dead/live cells. For quantification of proliferating Gfi1-YFP^+^ cells, the cells were stained with Ki67 antibody, followed by the appropriate Alexa Fluor 594 (Jackson Immuno) secondary antibody, or incubated with Click-iT reaction (Invitrogen) for EdU detection. Gfi1-YFP^+^ cells obtained from the intestinal crypts and stored in Trizol (Invitrogen) for RNA analysis. Primary and secondary antibodies are listed in Table 2.

### Real-Time PCR

Cells were sorted from intestinal crypts and RNA was extracted using a Qiagen RNeasy mini kit. A total of 1 μg RNA was used to synthesize complementary DNA using Superscript III First Strand Synthesis System (Invitrogen) following the manufacturer’s instructions. Quantitative PCR was performed with Brilliant III Ultra Fast SYBR Green Master Mix (Agilent Technologies) using the primers listed in Table 1.

**Table 1.**
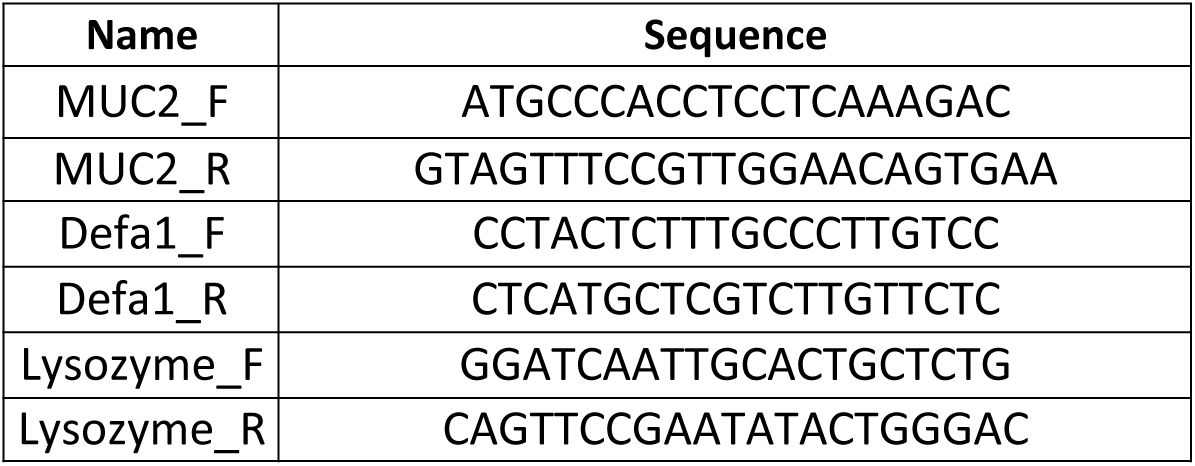
qPCR primers.

**Table 2.**
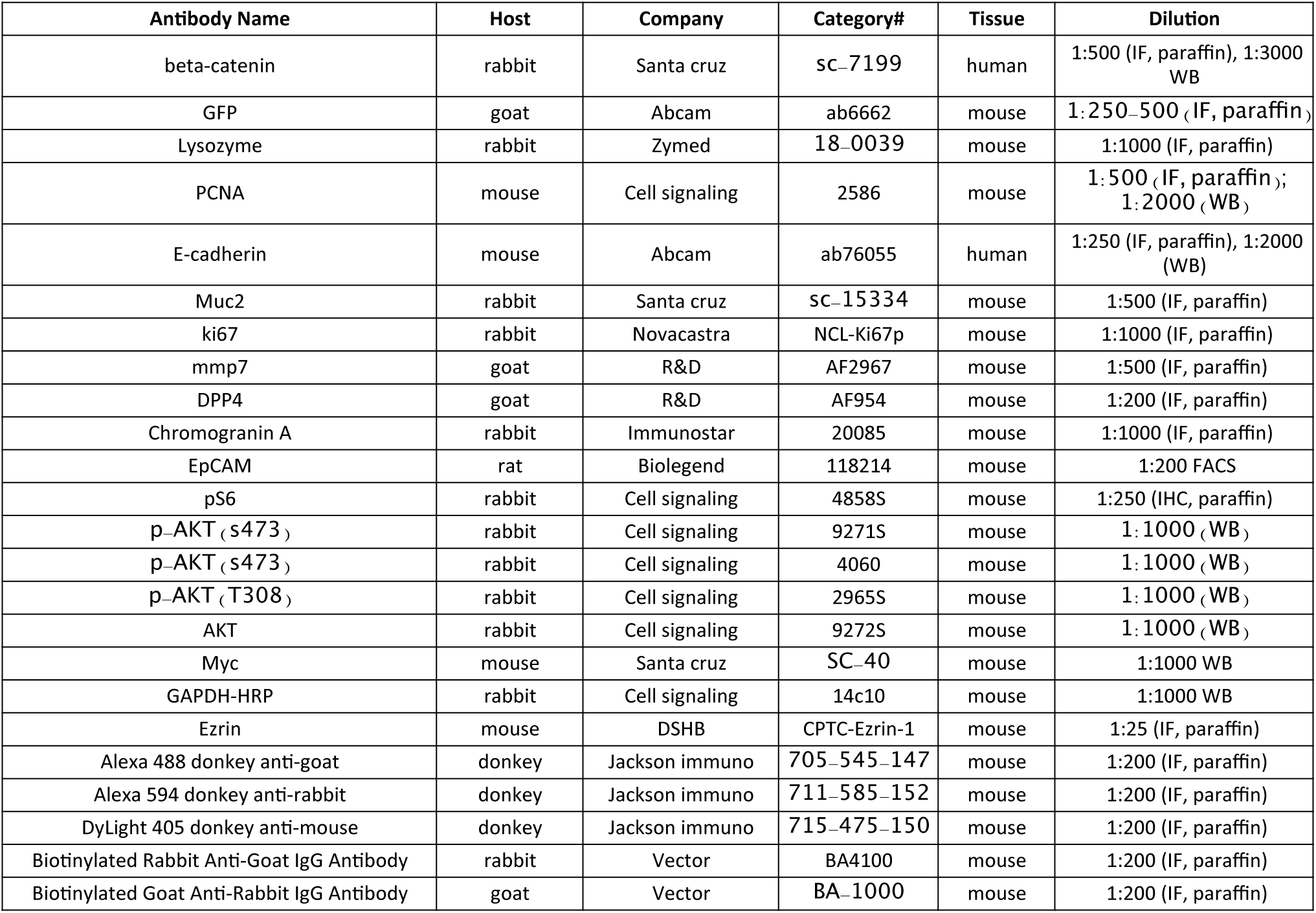
Antibody list.

## Results

### Gfi1 marks mature Paneth and goblet cells in the homeostasis

Gfi1 is a transcription factor that regulates secretory cell lineages. To investigate Gfi1^+^ secretory cells, we generated Gfi1^cre^ knock-in mice crossed to the Cre reporter strains R26^LSL-GFP^, R26^LSL-YFP^ or R26^LSL-tdtomato^ (Figure 1A). Immunofluorescent staining showed Gfi1 lineage cells are co-localized with the Paneth cell marker, lysozyme (Lyz) and the goblet cell marker, mucin 2 (Muc2) in the small intestine (Figure 1B). Quantification revealed that over 93% Lyz^+^ Paneth cells are Gfi1-YFP cells, and over 99% Muc2^+^ goblet cells are Gfi1-YFP cells (Figure 1C) in the small intestine. It is important to note that Gfi1 is also expressed in immune cells, which we confirmed demonstrating that Gfi1-YFP cells in the lamina propria are CD45^+^ immune cells (Supplemental Figure 1). We further demonstrated that FACS-isolated Gfi1-YFP^+^ cells were enriched in the markers *Lyz* and *Muc2*. On the other hand, YFP negative cells did not express these Paneth and goblet cell markers (Figure 1D). We concluded that Gfi1 lineage cells are differentiated Paneth and goblet cells in the small intestine.

**Figure 1.**
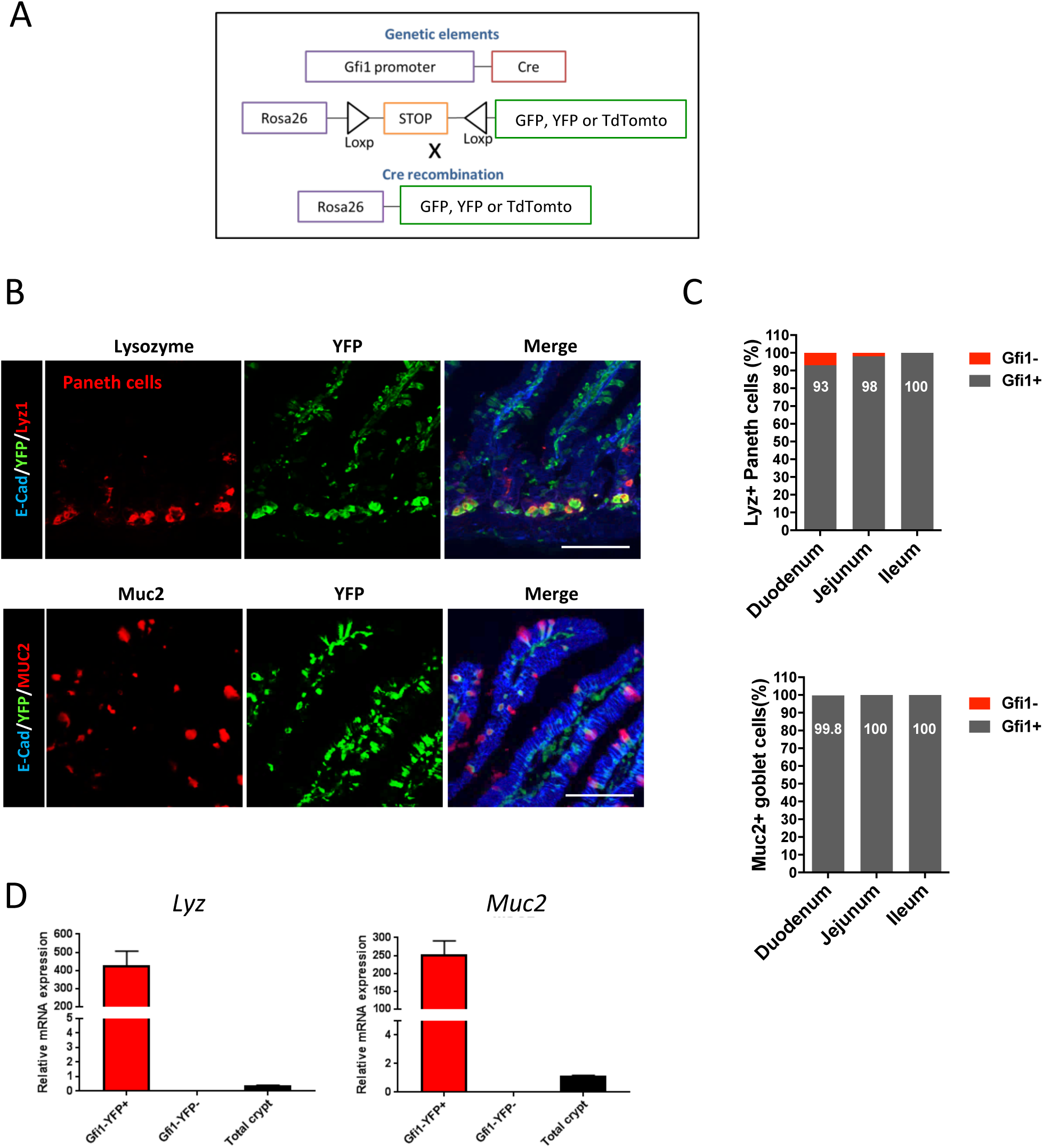
Gfi1^+^ lineage cells express Paneth and goblet cell markers. (A) An schematic representation of the cross between the Gfi1Cre Knock-in mouse and the Cre-mediated reporter *R26-GFP, R26-YFP or R26-tdTomoto to* generate the Gfi1 reporter mouse. (B) Immunostaining showed Gfi1-YFP cells from *Gfi1^cre/+;^ R26^LSL-YFP^ mouse* were co-localized with Paneth cell marker, lysozyme (Lyz) (top panel) and goblet cell marker, mucin 2 (Muc2) (bottom panel). Green: YFP; Red: Lyz (top), Muc2 (bottom); Blue: E-cadherin. Scale bars: 100 μm. (C) Quantification of Gfi1-YFP cells in Lyz+ Paneth cells and Muc2+ goblet cells. (D) Gfi1-YFP cells were sorted by FACS and analyzed for *Lyz* and *Muc2* expression by qPCR.

### Gfi1 lineage cells re-enter cell cycle upon crypt damage

To examine the effects of chemotherapy-induced injury on Lgr5^+^ and Gfi1^+^ lineage cells, we treated *Lgr5*-eGFP-IRES-CreERT2 and Gfi1^cre/+;^ R26^LSL-YFP^ mice with Doxorubicin (20 mg/kg, i.p.) to induce intestinal injury. By 48 hours post-injury, histological staining showed mucosal injury with crypt loss and shortened villi (Figure 2A). However, the total number of crypts was restored to baseline 7 days after injury, indicating that doxorubicin induced an acute and reversible intestinal injury (Figure 2B). We found a significantly decreased number of Lgr5 stem cells after doxorubicin treatment, reaching a nadir at 48 hours-post treatment (Figure 2C and 2D). Interestingly, Gfi1 lineage cells persisted and expanded in the crypts following doxorubicin injury. In uninjured mice, Gfi1-lineage cells were post-mitotic, as indicated by immunofluorescent staining and flow cytometry quantifying the fraction of Gfi1-lineage epithelial cells that were marked by the proliferative marker, Ki67 (Figure 2 E-F). However, by 24–48 hours following injury, Gfi1-lineage cells re-entered the cell cycle. We confirmed this by co-staining Gfi1-lineage cells with Ki67 and the S-phase marker EdU 48 hours after doxorubicin treatment; EdU+/Ki67+/Gfi1-YFP+ cells were rare in uninjured mice (Figure 2 F).

**Figure 2.**
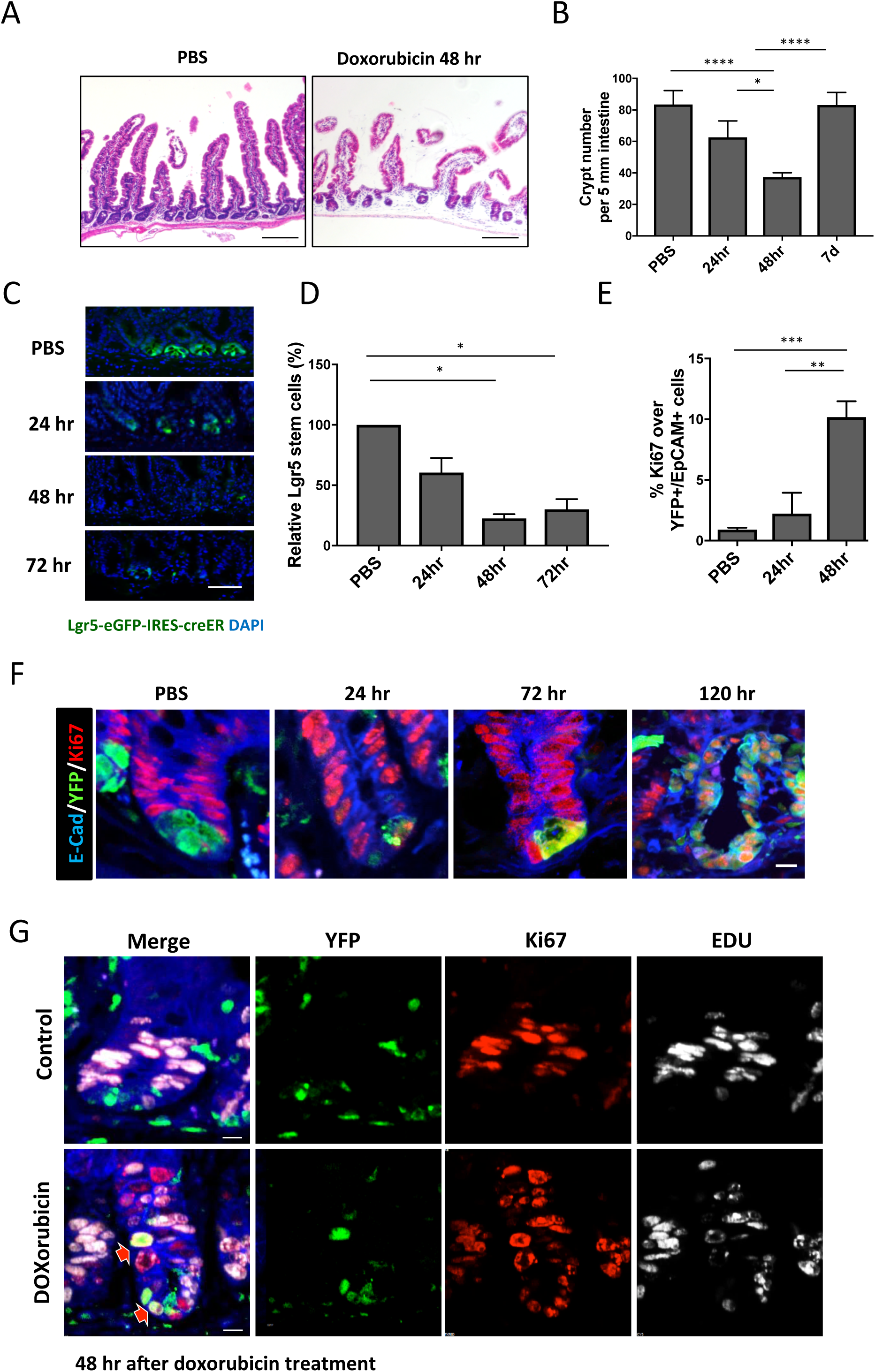
Gfi1 lineage cells re-enter cell cycle upon stem cell depletion. (A) Histological staining by H&E showed intestinal mucosal damage in mouse 2 days after doxorubicin treatment. (B) Quantification of crypt number in control and doxorubicin-treated mice. (C) Lgr5-EGFP mice were treated with doxorubicin and intestinal tissues were collected. Lgr5^+^ stem cell were analyzed by immunofluorescent staining. Green: GFP; Blue: DAPI. Scale bar, 50 μm. (D) Quantification of Lgr5^+^ cells by FACS. To determine whether Gfi1+ secretory cells re-enter cell cycle, Gfi1^cre/^+^;^ R26^LSL-YFP^ mice were treated doxorubicin for 48 hr. Ki67 staining (F) or EdU staining (G) were used to label cycling cells. Arrows indicated proliferative Gfi1-YFP cells (YFP+Ki67+EdU+). (E) Ki67+YFP+ live epithelial cells (EpCam+) were quantified by FACS. Blue: E-cadherin; Green: YFP; Red: Ki67; White: Edu. Scale bar, 10 μm.

### Gfi1^+^ secretory cells display plasticity following tissue injury

By 7 days after injury, Gfi1-lineage cells formed contiguous crypt-villus axis "ribbons" of cells (Figure 3A), suggesting these cells were functioning as multipotent stem cells. We had previously never observed such Gfi1-lineage ribbon tracing events in control mice without injury. Marker analysis using immunofluorescent staining demonstrated that Gfi1-lineage ribbon tracing included Paneth cells, goblet cells, enteroendocrine cells and enterocytes after injury (Figure 3B). To further demonstrate the conversion of Gfi1-lineage cells to CBC stem cells, we generated Lgr5-eGFP-IRES-CreERT2; Gfi1^cre/+;^ R26^LSL-tdtomato^ mice to label both Lgr5^+^ stem cells (GFP+) and Gfi1-lineage cells (Tomato+). In uninjured control mice, we never observed co-expression of GFP with Tomato (Figure 3C); however, 48 hours after doxorubicin injury we could observe GFP^+^ CBC cells derived from the Tomato+ Gfi1 lineage (Figure 3C). Moreover, Gfi1-lineage cells were more resistant to doxorubicin-induced injury, as evidenced by negative staining for DNA damage marker γH2AX and the apoptotic marker caspase-3 (Figure 3D).

**Figure 3.**
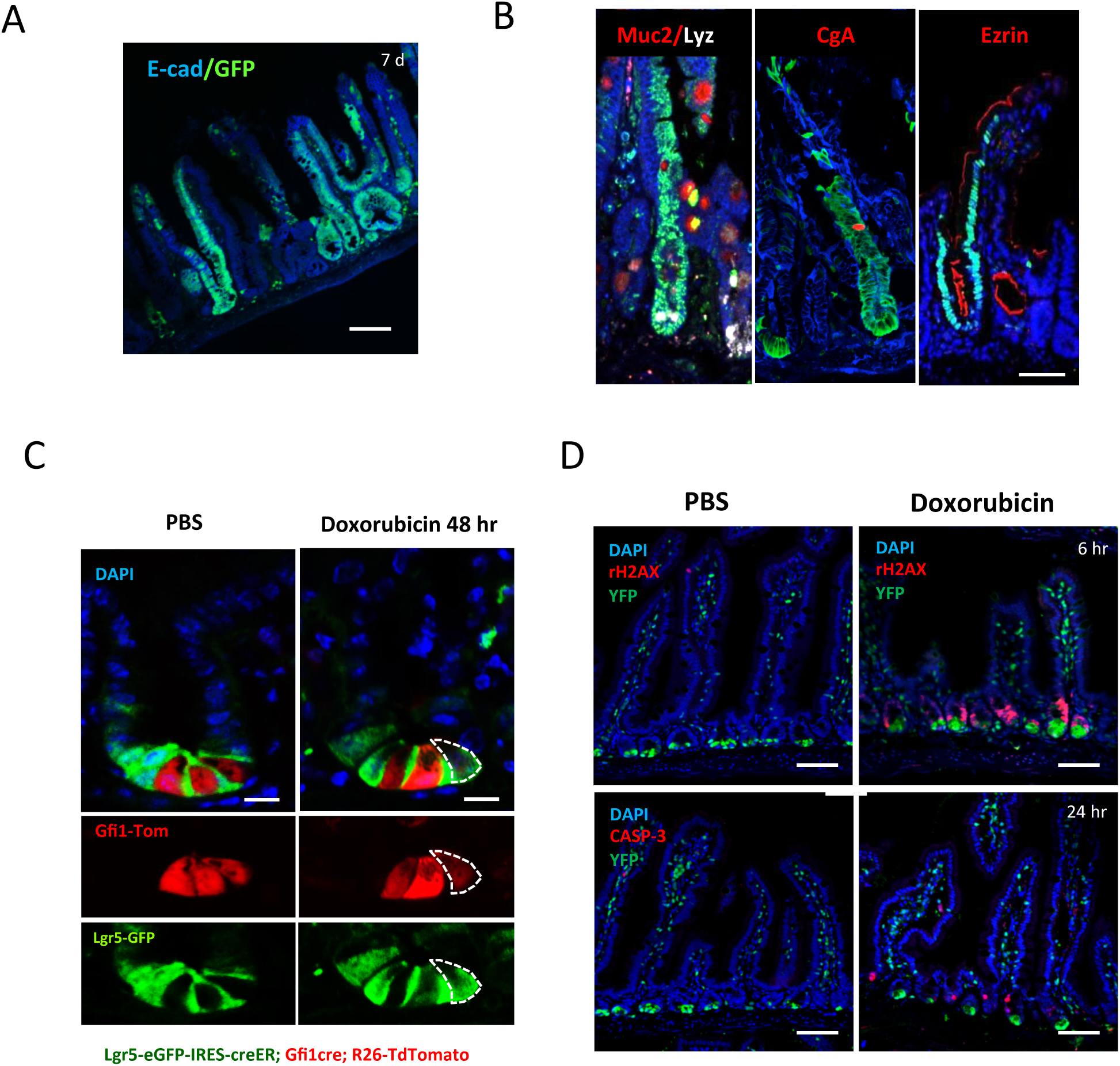
Gfi1 secretory cells revert to Lgr5^+^ stem cells and repopulate to all cell lineages. (A) Histological analysis of GFP staining of the intestine in *Gfi1^cre/+^*^;^ *R26^LSL^*^−^*^GFP^* mouse 7 days after doxorubicin treatment. Lineage tracing revealed that the formation of numerous GFP^+^ stem cell ribbons. (B) Gfi1-GFP^+^ ISCs are multipotent by immunofluorescenct antibody detection of lineage markers within Gfi1-lineage ribbons including Muc2 for goblet cells, Lyz for Paneth cells, chromogranin A (CgA) for enteroendocrine cells and Ezrin for enterocytes. (C) *Lgr5-*EGFP; *Gfi1^cre/+^*; *R26^LSL^*^−^*^YFP^* mouse was treated with doxorubicin for 48 hours. Fluorescent images showed Gfi1-YFP cell co-expressed Lgr5 stem cell marker. Dashed lines indicated GFP^+^Tomato^+^ cell. Scale bar: 10 μm. (D) Immunofluorescent analysis showed negative staining of DNA damage marker, rH2AX and apoptotic marker, caspase 3(CASP-3) after doxorubicin-induced injury. Scale bare: 100 μm.

### Reversion of Gfi1 secretory cells in different injury models is dependent on stem cell depletion

We next sought to determine whether Gfi1-lineage cells could contribute to the stem cell pool following other types of experimentally-induced intestinal injuries. We compared 3 different injury models: whole-body irradiation, immune-induced injury by systemic activation of the T-cell receptor with anti-CD3, and rotavirus (RV) infection (Figure 4). Irradiation and anti-CD3 treatment have previously been shown to cause stem cell damage ^36, 41^. On the other hand, RV infects cells via the villus tips, and does not infect or damage active and reserve stem cells ^37^. We observed Gfi1-lineage ribbon tracing events following either 10 Gy total body irradiation or anti-CD3 treatment, but not in the RV infected mice, indicating that stem cell depletion acts as a driving force of Gfi1^+^ cell reversion. The extent of reversion was generally correlated with the severity of injury, as assessed by dysmorphic crypts. We also treated Gfi1-null mice (Gfi1^cre/cre^; R26^LSL-GFP^) with doxorubicin. Interestingly, we found that Gfi1^+^ lineage tracing was still present in Gfi1-null mice (Figure 4B), suggesting that Gfi1 protein may not play a critical role in stem cell reversion. Taken together, these data suggest that Gfi1^+^ lineage cells revert to Lgr5^+^ stem cells upon stem cell damage and repopulate to all intestinal lineages.

**Figure 4.**
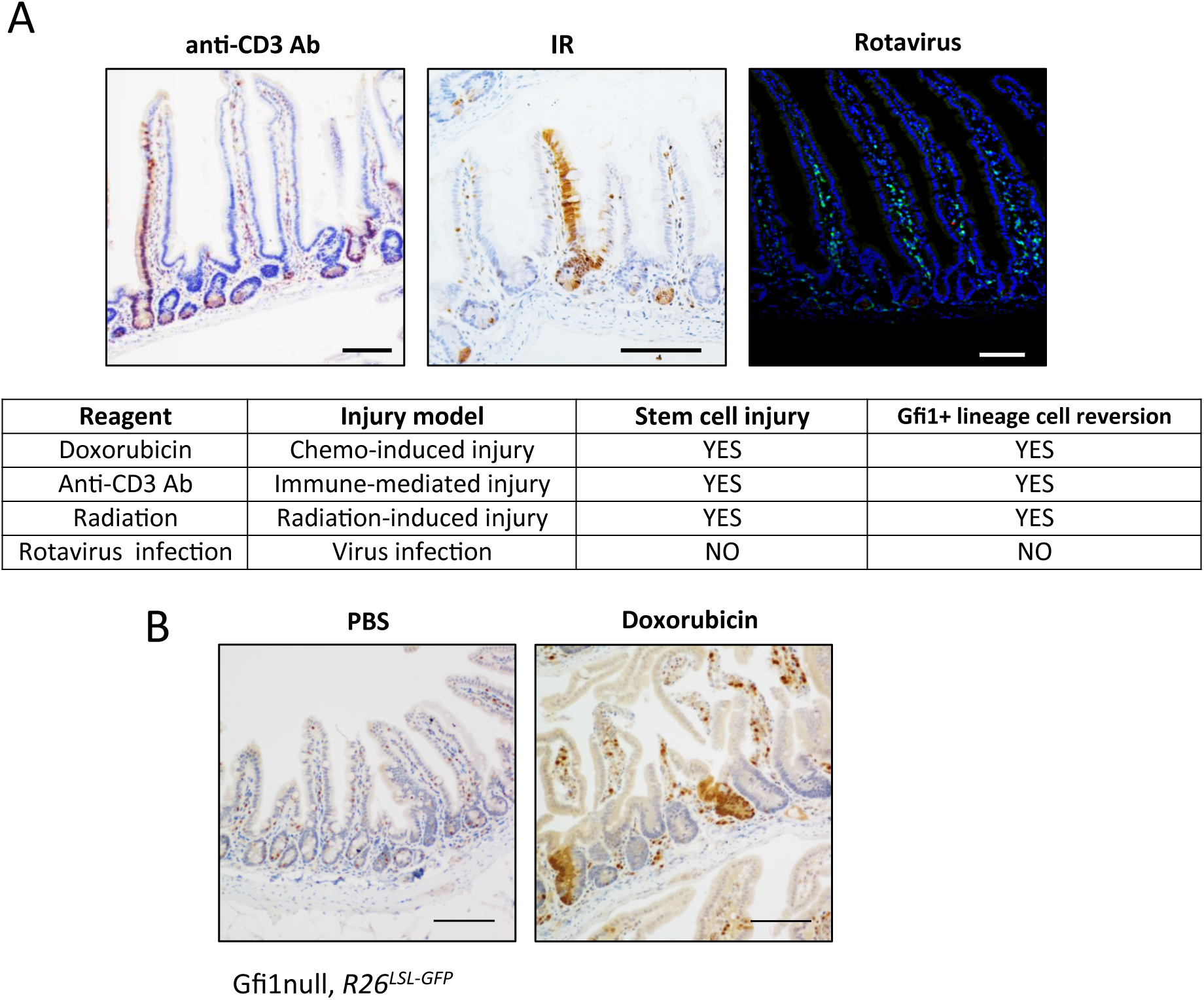
The extent of reversion was correlated with the severity of injury. (A) *Gfi1^cre/+^; R26^lsl^*^−^*^yfp^* mice were treated with anti-CD3 ab (left panel), 10 Gy radiation (middle panel) and rotavirus (right panel). By 7 days after treatment, we analyzed Gfi1-lineage tracing events by YFP reporter. A summary of this experiment is shown in Table. To examine whether Gfi1 protein function is required for stem cell reversion. We treated Gfi1null mice (Gfi1^cre/cre^; *R26^LSL-GFP^*) mice with doxorubicin for 7 days. Immunostaining showed Gfi1-lineage traces in doxorubicin-treated mouse but not in PBS-treated mouse. Scale bare: 100 μm.

### Establishment of *in vitro* injury model in mouse enteroids

We found that the expression of phospho-S6 (pS6) Ribosomal Protein, a downstream target of the AKT signaling pathway, was persistently upregulated in the small intestine for 7 days after doxorubicin-induced injury (Figure 5A). To better understand the mechanism underlying stem cell reversion, we established an *in vitro* doxorubicin-induced injury model using enteroids. Enteroids derived from Gfi1^cre/+;^ R26^LSL-tdTomato^ mice showed that Gfi1-lineage cells were non-cycling (Ki67 negative) Lyz^+^ Paneth cells and Muc2^+^ goblet cells (Figure 5B). Consistent with the observation *in vivo*, immunoblots suggested that the expression of phospho-AKT (pAKT) and pS6 was upregulated following doxorubicin treatment (Figure 5C). The activation of the AKT signaling pathway could be further manipulated by either PTEN or AKT inhibitors (Figure 5C). These results indicated that intestinal enteroids exhibited similar response to doxorubicin *in vivo*, and could be used as an *in vitro* injury model.

**Figure 5.**
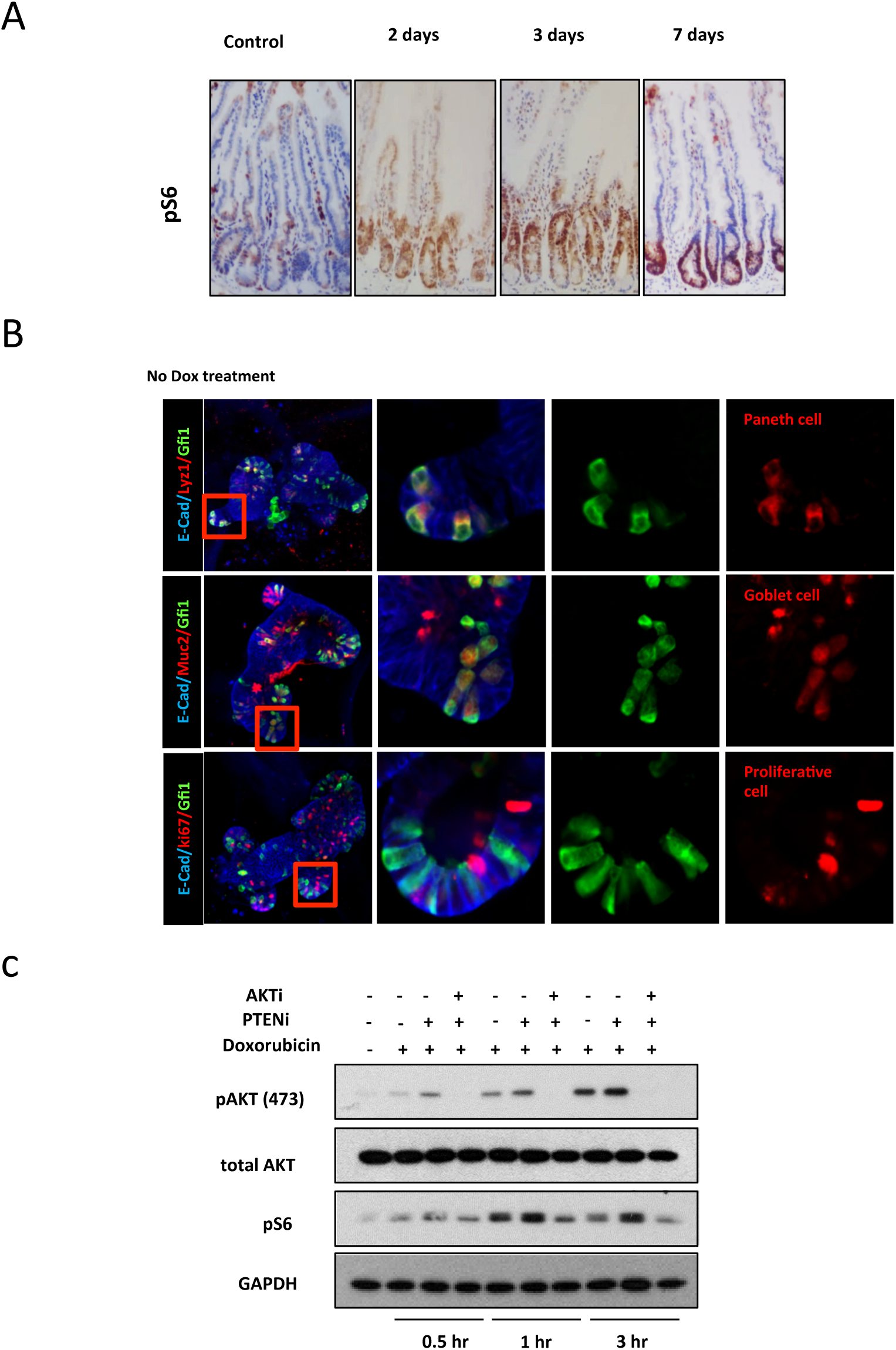
AKT/mTOR signaling is upregulated in injured tissue. (A) Immunostaining by pS6 antibody showed the induction of pS6 expression that was persisted for 7 days after injury. (B) We generated enteroids from *Gfi1^cre/+^; R26^LSL^*^−^*^YFP^* mice and analyzed cells by secretory cell marker, Lyz and Muc2 and proliferative marker, Ki67. (C) Immunoblotting showed the expression of pAKT, total AKT, pS6 and GAPDH after doxorubicin, PTEN inhibitor or AKT inhibitor treatment.

### Activation of PI3K/AKT and Wnt signaling pathways enhances cell survival and stem cell reversion following injury

In our *in vitro* injury model, we examined the effects of various inhibitors and growth factors on cell growth and Gfi1^+^ stem cell reversion following injury (Figure 6A). After a small screen, we demonstrated that the PTEN inhibitor (PTENinh), when combined with high R-spondin (H-Rspo) treatment for 48 hours immediately after the induced injury, significantly improved survival of enteroids *in vitro* (Figure 6B and C). Immunoblots confirmed the activation of pAKT after PTENinh treatment (Figure 5C, Supplementary Figure 2B) and upregulation of Myc, a Wnt downstream target after high R-spondin treatment (Supplementary Figure 2A). Moreover, we were able to identify reversion of Gfi1^+^ cells after injury. Initially, a cluster of Gfi1 lineage cells was observed in a subset of enteroids after doxorubicin treatment (Supplementary Figure 3). After passaging, we were able to find several enteroids that expressed the Gfi1-lineage marker homogeneously throughout the enteroid (Supplementary Figure 3), indicating that Gfi1^+^ cells had reverted to stem cells after injury and produced all of the cells within the passaged enteroid. Most importantly, we found that PTENinh and H-Rspo treatment significantly increased number of Gfi1-lineage derived, revertant enteroids following injury (Figure 6D&E), indicating that PI3K/AKT and Wnt stimulation enhanced Gfi1-lineage cell reversion. We further showed that the AKT inhibitor blocked PI3K/AKT and Wnt-induced enteroid growth and Gfi1-lineage cell reversion, indicating that AKT signaling plays a critical role in the regeneration and cellular reversion process (Figure 6C&E).

**Figure 6.**
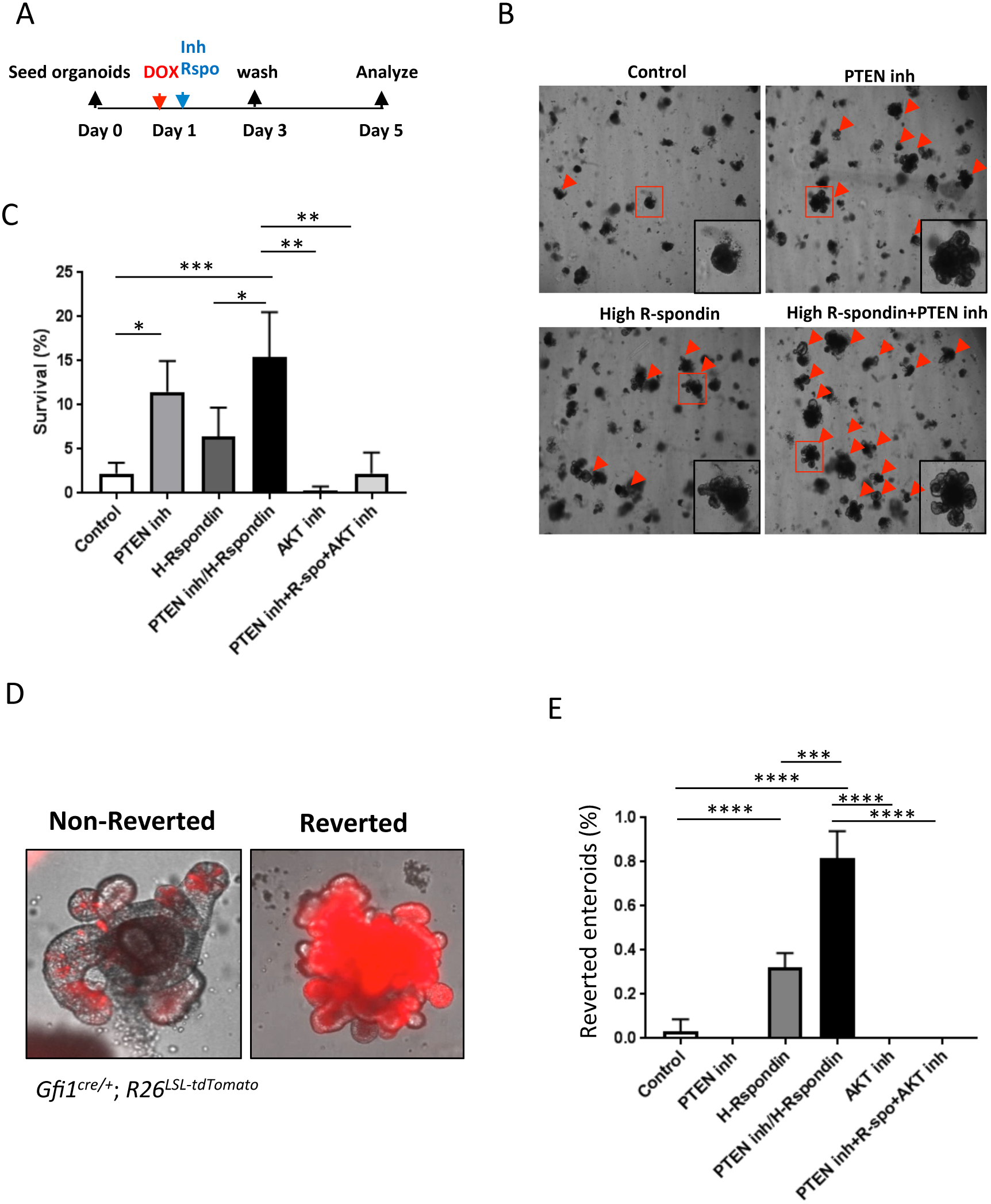
PI3/AKT and Wnt stimulation enhance cell survival and stem cell reversion following injury *in vitro*. (A) Scheme of doxorubicin, inhibitors and growth factors treatment in enteroids. (B) Bright field images showed the enteroid growth and budding in enteroids after treatment. Arrows indicated live enteroids. (C) Quantification of enteroid survival in different treatment groups. (D) Images demonstrated non-reverted versus reverted enteroids with or without injury. (E) Quantification of reverted enteroids in different treatment groups.

### Wnt and PI3K/AKT signaling improve tissue regeneration *in vivo*

Previous studies indicated Wnt stimulation by Ad-Rspo1 induced crypt hyperplasia and when combined with Slit2, there was reduced stem cell loss after injury ^30^. Furthermore, this treatment paradigm provided significant chemoradioprotection. To determine whether increased PI3K/AKT and Wnt enhanced tissue regeneration *in vivo*, we treated *Gfi1^cre/+;^R26*^−^*^LSL^*^−^*^YFP^* mice with Ad-Rspo1 or Ad-Fc encoding a control immunoglobulin lgG2a Fc fragment 2 days before doxorubicin administration. An additional cohort of mice were treated with PTENinh after injury, or a combination of Ad-Rspo1 pre-treatment and PTENinh treatment 1 hour after doxorubicin administration. Intestinal tissues were collected 7 days after doxorubicin treatment. Histological staining showed a significant increase in the crypt depth in the Ad-Rspo1-treated mice compared with the mice receiving the control Ad-Fc treatment (Figure 7A). Crypt number and depth were significantly increased in the Ad-Rspo1- and PTENinh-treated mice (Figure 7C and 7D). Although we did not observe an increase in the number of Gfi1-lineage ribbon tracing events in the double-treated mice (Figure 7E), we found that the number of crypts marked in revertant clones, and presumably derived from single reversion events, was significantly increased in Rspo1/PTENinh-treated mice (Figure 7B, F). Taken together, our data suggests that administering Ad-Rspo1 followed by treatment with PTENinh enhanced tissue regeneration *in vivo*.

**Figure 7.**
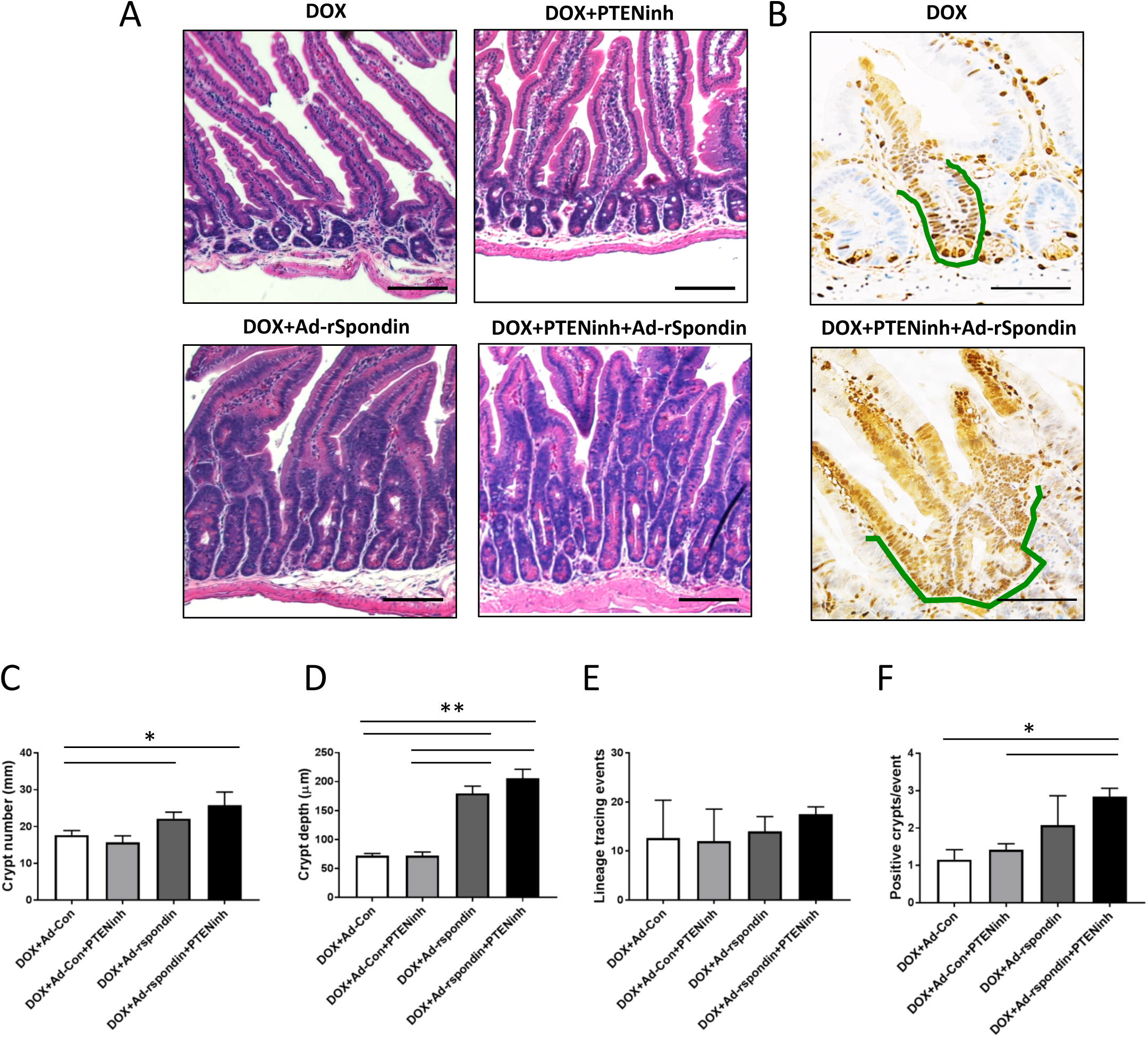
Activation of PI3K/AKT and Wnt improves tissue regeneration *in vivo*. (A) Histological analysis by H&E in *Gfi1^cre/+^; R26^LSL^*^−^*^YFP^* mice with different treatments. (B) We analyzed Gfi1-lineage tracing events using immunostaining by GFP antibody. Crypt number (C), crypt depth (D), lineage tracing events (E) and number of positive YFP+ crypts per event (F) were quantified. Green lines indicated Gfi1-lineage tracing events.

## Discussion

Understanding the differences between active and injury-inducible ISCs’ response to damage has the potential to illuminate pathways to new therapeutic approaches for the selective protection of normal cells and to mitigate chemotherapy- or radiation-induced intestinal injury. Active Lgr5^+^CBC stem cells are continually in the cell cycle thus rendering them sensitive to radiation or chemotherapy. Previously, lineage tracing experiments revealed that reserve ISCs labeled by Bmi1, mTert, Hopx, Lrig1 and Sox9, display plasticity when active ISCs were depleted by injury ^6, 7, 9–12^. Recent studies suggested that secretory and absorptive progenitors reacquired stemness and act as potential stem cells under injury settings ^13–15^. Experimental efforts using single cell RNA-seq revealed that Bmi1 lineage cells are mature entroendocrine (EE) cells and express an entroendocrine cell signature ^16, 42^. Unlike other reporter mouse models which marked mostly early progenitors or cycling ISCs, our Gfi1 reporter mouse showed that in uninjured mice, Gfi1^+^ lineage cells are mature and non-cycling Paneth and goblet cells (Figure 1). After injury, Gfi1^+^ secretory cells are more resistant to chemo-induced DNA damage and apoptosis, whereas other crypt cells including Lgr5^+^ ISCs undergo extensive DNA damage and cell death (Figures 2 & 3). This injury triggers the dedifferentiation of Gfi1^+^ secretory cells, which is accompanied by cell cycle re-entry and reversion to replace the depleted ISC pool (Figures 2 & 3). Intestinal regeneration following injury has been studied using several experimental models. Severe injury caused by cytotoxic drugs, such as doxorubicin, or irradiation results in the transient depletion of ISCs. Similarly, acute inflammatory injury using anti-CD3 antibody to activate T cells results in apoptosis of active ISCs, which recover a few days later^36^. RV infection is another model to study the injury-repair responses. Previous studies demonstrated that RV only infected enterocytes on the villi, while ISCs within the crypt remained intact without suffering any damage^37^. Comparing the four different injury models, it appears that the depletion of active ISCs is required for Gfi1^+^ secretory cells to revert to Lgr5^+^ ISCs and repopulate all intestinal cell lineages (Figure 4).

It is of great clinical significance to study mechanisms involved in the maintenance of tissue homeostasis and regeneration. Wnt signaling is known to be important for the regulation of stem cell renewal, proliferation, and differentiation of intestinal epithelial cells ^43^. During tissue regeneration, Wnt signaling is necessary for promoting healing responses and driving the cell’s regenerative capacity^44^. R-spondin proteins act as synergistic activators of the Wnt signaling pathway that drive ISCs self-renewal and tissue regeneration. Previous work showed that elevating Wnt activity using Ad-Rspondin 1 prior to irradiation or chemotherapy, protects normal intestinal cells from injury ^29, 30^. PI3K-AKT-mTORC1 is another important regulatory signaling pathway for intestinal epithelial tissue homeostasis ^34^. Our results indicated that injury resulting from doxorubicin treatment induced pAKT expression in an *in vitro* enteroid culture system, and pS6 expression levels in both enteroids and after doxorubicin treatment in mice (Figure 5). These data collectively suggest that PI3K-AKT-mTORC1 signaling plays an important role in the regenerative process. Our enteroid data showed PTENinh or R-spondin 1 treatment alone improved enteroid regrowth following injury (Figure 6). Interestingly, we observed that extra R-spondin 1 increased the capacity for reversion of Gfi1^+^ cells to stem cells. In contrast, we did not see an increase in stem cell reversion when PI3K/AKT was induced by PTENinh. However, PTENinh combined with R-spondin 1 treatment significantly enhanced both enteroid survival and stem cell reversion, suggesting that there is crosstalk between the PI3K/AKT and the Wnt signaling pathways. Further, we showed that blocking AKT using an inhibitor eliminated all growth and reversion, suggesting that PI3K/AKT is epistatic to the Wnt signaling pathway with regard to survival and stem cell plasticity. Further studies are needed to more definitively elucidate the interaction between the PI3K/AKT and Wnt pathways in the injury-repair process. Additional studies using the enteroid system to explore the mechanisms and potential mitigators of stem cell injury are needed and will be facilitated by high-throughput screening methodologies as recently described ^45, 46^.

Our time course data showed that Gfi1-lineage ribbon tracing events appear to emerge from the crypt base (Figure 2F), indicating that Paneth cells are more likely to revert and contribute to injury-induced ISCs. However, our Gfi1 reporter mice marked both Paneth and goblet cells, so we cannot rule out the possibility that crypt-resident goblet cells may also revert to Lgr5^+^ stem cells following injury. There is an unlikely possibility that other cells, when reverting to injury-induce ISCs, turn on Gfi1 before fully reverting. Finally, our *in vivo* treatment showed that R-spondin 1 and PTENinh treatment enhanced Gfi1^+^ cells mitotic activity, suggesting that manipulation of the Wnt and PI3K/AKT pathways during tissue damage may enhance injury-induced ISCs regeneration and reconstitute lost tissue.

In summary, lineage tracing revealed that Gfi1^+^ cells retain cell fate plasticity and revert to Lgr5^+^ stem cells following severe injury. Our results demonstrate that terminally differentiated secretory cells, primarily Paneth cells, can re-enter the cell cycle and revert to Lgr5^+^ ISCs to repopulate the depleted ISC pool and facilitate intestinal regeneration. Furthermore, we find that the Wnt and PI3K/AKT signaling pathways are key players in tissue repair and regeneration by enhancing survival, stem cell reversion, and clonal expansion. These studies not only improve our current understanding of injury-induced ISCs that contribute to intestinal epithelial regeneration, but also offer insights for developing novel therapeutic strategies to mitigate and enhance regeneration after tissue injury.

## Acknowledgments

We thank Dr. Milton Finegold, Dr. Jason Heaney, Dr. Keith Chan, Dr. Scott Magness, and Dr. Cullen Taniguchi for critical input on experiments. We thank the members of the Estes lab at Baylor College of Medicine, Kuo lab at Stanford University, Taniguchi lab at MD Anderson for their technical assistance. We thank Dr. Richard Finnell for manuscript editing. We thank the Texas Medical Center Digestive Disease Center for their support with funding from the National Institutes of Health (P30DK56338), the Intestinal Stem Cell Consortium with funding from the NIH through grants (U01 DK103168, U01 DK103168-03S), NIH grants R01 CA142826 (N.F.S.) and F99 CA212433 (Y.H.L). We thank the Integrated Microscopy core at Baylor College of Medicine for their support with funding from the NIH (DK56338 and CA125123), the Cytometry and Cell Sorting Core at Baylor College of Medicine with funding from the National Institutes of Health (P30 AI036211, P30 CA125123, and S10 RR024574) and the Dan L. Duncan Cancer Center.

---

**Supplementary Figure 1.**
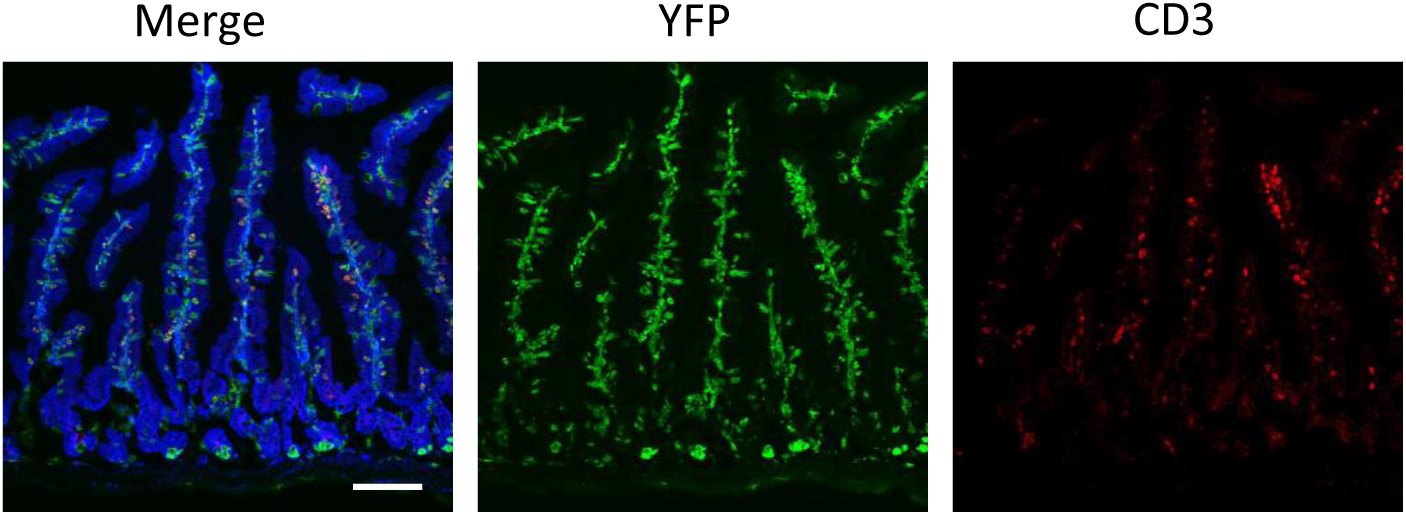
Immune cells express Gfi1 in the small intestine. (A) Immunofluorescent staining showed Gfi1-lineage cells in lamina propria in the small intestine are CD3+ immune cells. Blue: E-cadherin; Green: YFP; Red: CD3. Scale bar: 100 μm.

**Supplementary Figure 2.**
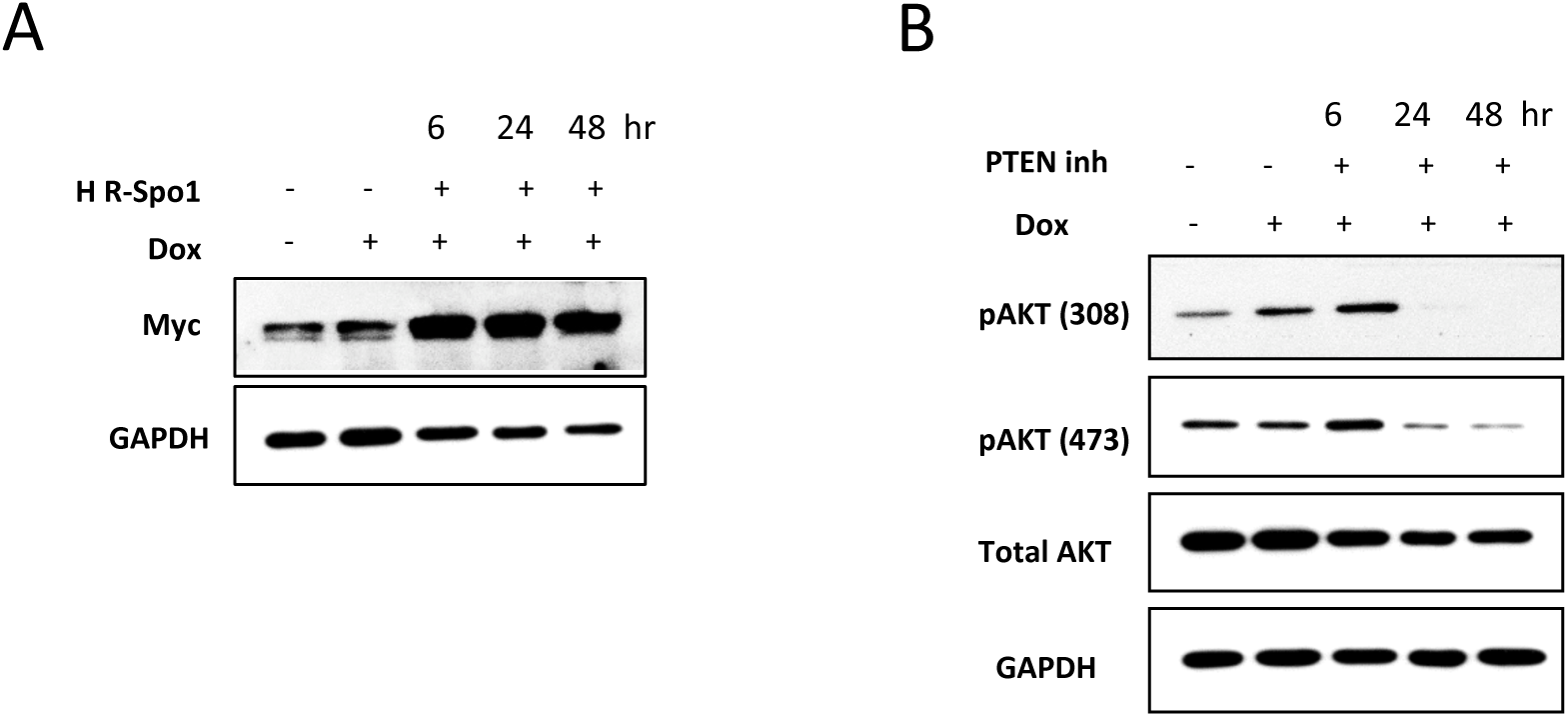
Induction of Wnt and AKT signaling pathways by high R-spodin1 and PTENinh treatment. (A) Enteroids were treated with doxorubicin for 4 hours and then high R-spondin1 (Rspo1) conditioned media (60%) (A) or PTENinh (B) were added into enteroids for 6, 24 and 48 hours. Myc, pAKT(Ser473), pAKT(Thr308) and total AKT were detected by Western blot. GAPDH was used as internal control.

**Supplementary Figure 3.**
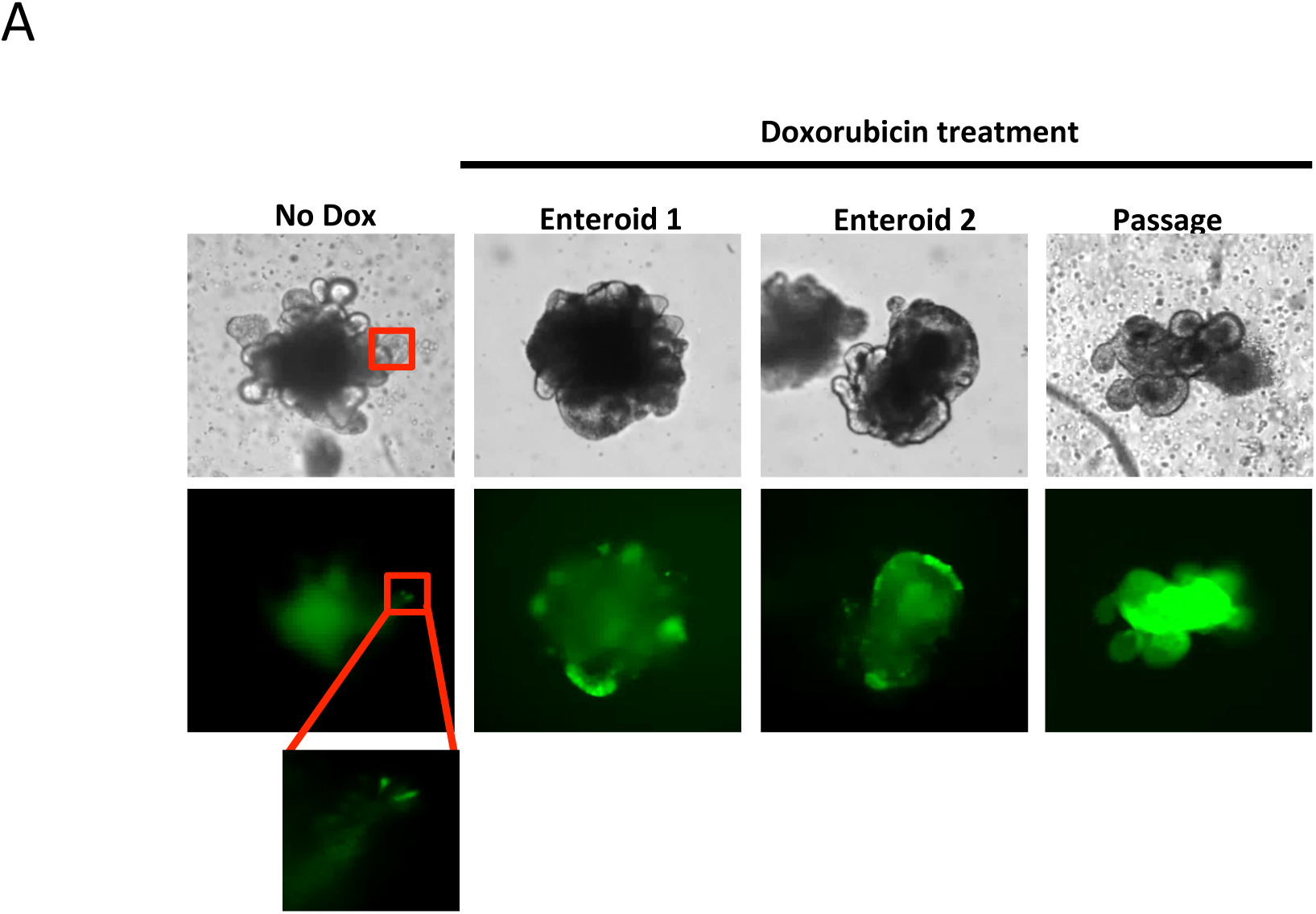
Gfi1^+^ secretory cells possess sternness after injury. Live images showed enteroids derived from *Gfi1^cre/+^; R26^LSL^*^−^*^YFP^* in uninjured or injured condition. By 12 days after doxorubicin treatment, a cluster of Gfi1-lineage cells was showed in enteroids. After passage, several YFP-homogeneous enteroids were showed in doxorubicin-treated group but not in control group.

